# *De novo* assembly and phasing of dikaryotic genomes from two isolates of *Puccinia coronata* f. sp. *avenae*, the causal agent of oat crown rust

**DOI:** 10.1101/179226

**Authors:** Marisa E. Miller, Ying Zhang, Vahid Omidvar, Jana Sperschneider, Benjamin Schwessinger, Castle Raley, Jonathan M. Palmer, Diana Garnica, Narayana Upadhyaya, John Rathjen, Jennifer M. Taylor, Robert F. Park, Peter N. Dodds, Cory D. Hirsch, Shahryar F. Kianian, Melania Figueroa

## Abstract

Oat crown rust, caused by the fungus *Puccinia coronata* f. sp. *avenae* (*Pca*), is a devastating disease that impacts worldwide oat production. For much of its life cycle, *Pca* is dikaryotic, with two separate haploid nuclei that may vary in virulence genotype, highlighting the importance of understanding haplotype diversity in this species. We generated highly contiguous *de novo* genome assemblies of two *Pca* isolates, 12SD80 and 12NC29, from long-read sequences. In total, we assembled 603 primary contigs for a total assembly length of 99.16 Mbp for 12SD80 and 777 primary contigs with a total length of 105.25 Mbp for 12NC29, and approximately 52% of each genome was assembled into alternate haplotypes. This revealed structural variation between haplotypes in each isolate equivalent to more than 2% of the genome size, in addition to about 260,000 and 380,000 heterozygous single-nucleotide polymorphisms in 12SD80 and 12NC29, respectively. Transcript-based annotation identified 26,796 and 28,801 coding sequences for isolates 12SD80 and 12NC29, respectively, including about 7,000 allele pairs in haplotype-phased regions. Furthermore, expression profiling revealed clusters of co-expressed secreted effector candidates, and the majority of orthologous effectors between isolates showed conservation of expression patterns. However, a small subset of orthologs showed divergence in expression, which may contribute to differences in virulence between 12SD80 and 12NC29. This study provides the first haplotype-phased reference genome for a dikaryotic rust fungus as a foundation for future studies into virulence mechanisms in *Pca*.

**Importance:** Disease management strategies for oat crown rust are challenged by the rapid evolution of *Puccinia coronata* f. sp. *avenae* (*Pca*), which renders resistance genes in oat varieties ineffective. Despite the economic importance of understanding *Pca*, resources to study the molecular mechanisms underpinning pathogenicity and emergence of new virulence traits are lacking. Such limitations are partly due to the obligate biotrophic lifestyle of *Pca* as well as the dikaryotic nature of the genome, features that are also shared with other important rust pathogens. This study reports the first release of a haplotype-phased genome assembly for a dikaryotic fungal species and demonstrates the amenability of using emerging technologies to investigate genetic diversity in populations of *Pca*.

## Introduction

Cultivated oat (*Avena sativa*) ranks sixth in global production among cereals like maize, rice, and wheat (1). In recent years, the demonstrated health benefits of oats and its expanded commercial applications have increased demand for the crop (2). Crown rust, caused by the pathogenic fungus *Puccinia coronata* f. sp. *avenae* (*Pca*), is the most devastating disease affecting production in nearly every oat growing region worldwide (2, 3) with yield losses due to infection reaching 50% (4).

*Pca* is a macrocyclic and heteroecious rust fungus (Puccinales, Basidiomycota) (2). Asexual or clonal reproduction of *Pca* occurs in oat, and its wild relatives, and involves repeated infection cycles mediated by urediniospores, which can perpetuate infection indefinitely (2). The infection process involves germination of urediniospores on the leaf surface, appressorium and penetration peg differentiation to allow host entry through a stomate, formation of a substomatal vesicle and the establishment of a colony by hyphal proliferation, and finally sporulation to produce more urediniospores. During infection, the fungus also forms haustoria, specialized feeding structures that allow nutrient uptake and secretion of effector proteins into the host cells (5). During the asexual cycle, *Pca* is dikaryotic, with each urediniospore containing two haploid nuclei, while the sexual cycle involves meiosis and infection of an alternate host of the genus *Rhamnus* (e.g. common buckthorn) by haploid spores and subsequent gamete fusion to re-establish the dikaryotic stage (2). Thus, the sexual cycle contributes to oat crown rust outbreaks both by generating an additional source of inoculum and by re-assorting genetic variation in the pathogen population.

Disease management strategies for oat crown rust rely heavily on breeding for race-specific resistance (2). However, *Pca* rapidly evolves virulence to new resistance genes and field populations are highly polymorphic with high numbers of races (pathotypes), which limits the efficacy of this approach (6). Resistance to *Pca* in *Avena* spp. conforms to the classical gene-for-gene model (7, 8), and is conditioned by dominant resistance (*R*) genes, which mediate recognition of cognate avirulence (*Avr*) factors in the pathogen. Plant *R* genes typically encode intracellular nucleotide binding and leucine-rich repeat (NLR) receptor proteins, which detect specific pathogen effector proteins and induce a localized hypersensitive response (9, 10). Evolution of new virulence traits occur due to changes in effector genes that allow the pathogen to escape recognition (11). Several *Avr* genes identified in the model flax rust, *Melampsora lini*, encode secreted proteins expressed in haustoria that are recognized inside host cells (12, 13). However, no *Avr* genes have been identified in *Pca* and the biological mechanisms generating genetic variability in *Pca* are unknown. Since *Pca* is dikaryotic, a virulence phenotype requires the loss of avirulence function of both alleles at the effector locus and thus emergence of virulence strains can be enhanced by sexual recombination. Nevertheless, the high diversity of virulence phenotypes in asexual populations suggests that additional molecular mechanisms like high mutational rates, somatic hybridization and somatic recombination play roles in generating variability in *Pca* (14-16).

Given their biotrophic lifestyle, most rust fungi are recalcitrant to *in vitro* culturing and genetic transformation, which hinders molecular studies of pathogenicity. Nevertheless, genome sequencing of a few rust species has provided insights into the biology and adaptations associated with parasitic growth (17-24). These resources have enabled the prediction of effector candidates and, in some instances, identification of *Avr* genes (13, 25). However, the large genome sizes of rust fungi sequenced to date (90-200 Mbp) compared to other pathogenic fungi (26-29), and high repetitive DNA content (over 50%) hamper *de novo* genome assembly from short-read sequencing, which leads to high fragmentation, mis-assembly errors and merging of two distinct haplotype sequences. The dikaryotic nature of rust fungi also means that current genome assemblies represent collapsed mosaics of sequences derived from both haplotypes and do not account for structural variation between haplotypes. Single-molecule real time (SMRT) sequencing has emerged as a powerful technology to achieve high-contiguity assembly of even repeat-rich genomes (30) and recently released algorithms enable the resolution of haplotypes in diploid genomes (31).

Here, we document the assembly of draft genome sequences for two *Pca* isolates with contrasting virulence phenotypes using SMRT sequencing and the FALCON assembler and FALCON-Unzip for haplotype resolution (31). The contiguity of the *Pca* assemblies is greatly improved compared to previous short-read *de novo* assemblies of rust species (20-22). We separately assembled the two haplotypes for over 50% of the haploid genome of each isolate. This revealed many structural differences between haplotypes and isolates, including large insertions/deletions covering both intergenic and coding regions. The *Pca* genomes were annotated utilizing expression data from different tissue types and life stages and a catalog of predicted secreted effectors was generated. To our knowledge, this study provides the first report of genome-wide haplotype resolution of dikaryotic rust fungi and the foundation to investigate the evolution of virulence factors and the contribution of haplotype variation to the pathogenicity of *Pca*.

## Results and discussion

### *Puccinia coronata* f. sp. *avenae* (*Pca*) isolates 12SD80 and 12NC29 show distinct virulence profiles

To build comprehensive genomic resources for virulence studies in *Pca* we selected two isolates, 12NC29 and 12SD80, from the 2012 USDA-ARS annual rust survey that show contrasting virulence profiles on an oat differential set (**Figure 1A** and **B**). Isolate 12SD80 is virulent on a broader range of oat differentials than isolate 12NC29, although recently released *Pc* resistance genes (*Pc91, Pc94, Pc96*) are effective against both isolates. Despite the different virulence profiles on specific *Pc* genes, both isolates showed similar infection progression over a seven-day time course on the susceptible oat variety Marvelous (**Figure 1C**). More than 90% of urediniospores germinated of which more than 60% differentiated an appressorium (penetration structure) in the first 24 hours of infection. Established colonies and first signs of sporulation were detected by 5 days post infection (dpi) and 40-50% of infection sites displayed sporulation by 7 dpi. Thus, both *Pca* isolates were equally aggressive in the absence of effective *Pc* genes.

**Figure 1.**
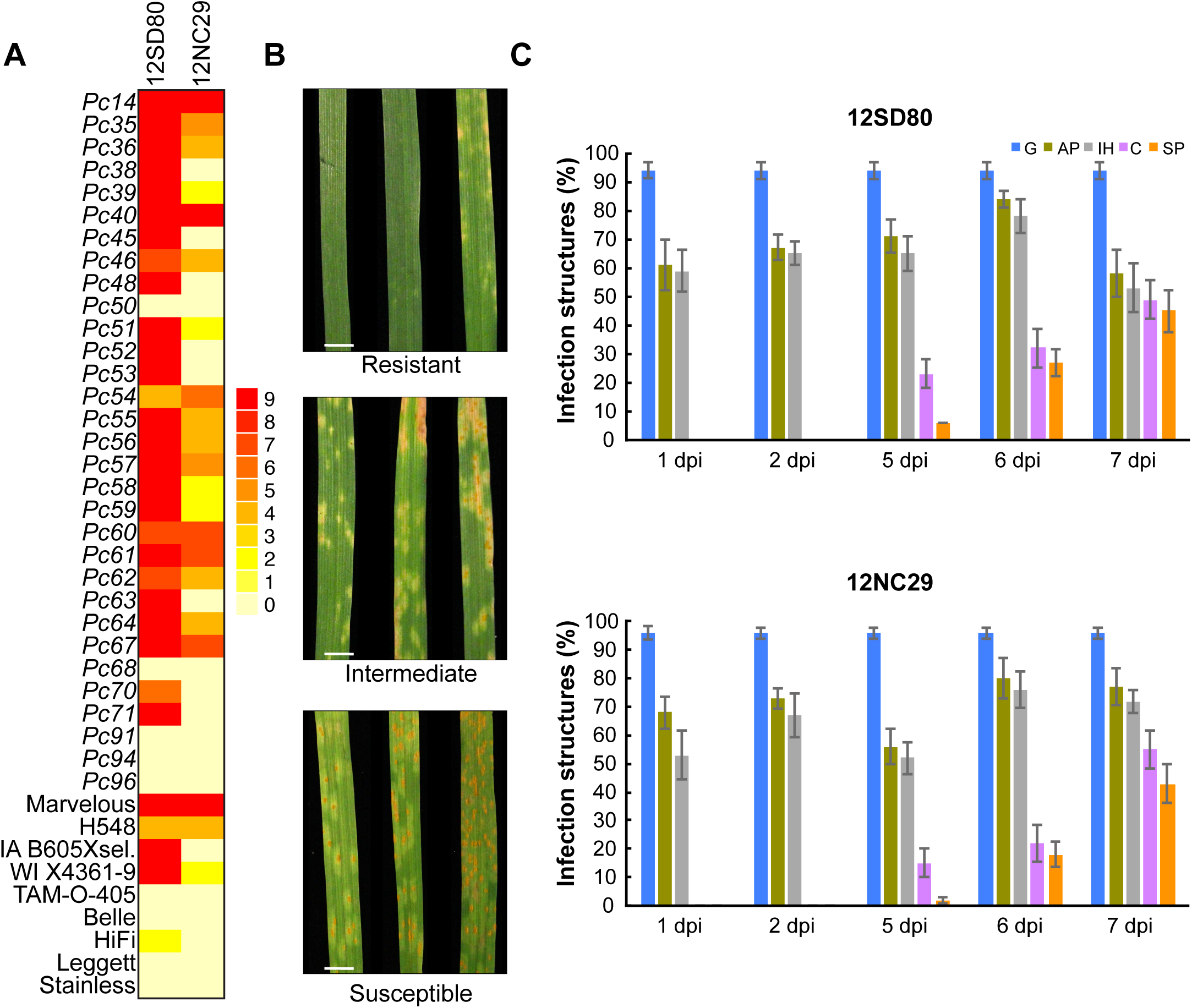
Phenotypic variation of *Pca* isolate virulence and colonization patterns in susceptible oat. (**A**) Heatmap showing virulence profiles of 12SD80 and 12NC29 on a set of 40 oat differential lines. (**B**) Photographs represent examples of infection types corresponding to full resistance or intermediate resistance, as well as susceptibility. Scale bar = 0.5 cm. (**C**) Quantification of infection structures of *Pca* isolates in the susceptible oat line Marvelous at 1, 2, 5, 6, and 7 dpi. Graphs show the percentage of urediniospores that have germinated (G), percentage of germinated spores which formed appressoria (AP), substomatal vesicles or primary infection hyphae (IH), established colonies (C), and sporulating colonies (SP). Error bars represent standard errors of three independent replicates.

### *De novo* genome assembly and haplotype-phasing of *Pca* isolates

High molecular weight DNA (>50 kbp) was extracted from germinated urediniospores of 12SD80 and 12NC29, and long-read sequence data was generated using SMRT sequencing. This yielded approximately 20.9 and 25.9 Gbp of filtered subreads for 12SD80 and 12NC29, respectively. The mean and N50 subread lengths were 6,389 and 8,445 bp, respectively, for 12SD80, and 6,481 and 8,609 bp for 12NC29 (**Table S1** and **Figure S1**). Subread distributions for both isolates extended to approximately 30,000 bp (**Figure S1**). Illumina sequencing was performed on the same samples and yielded approximately 6 and 7 Gbp of sequence information for 12SD80 and 12NC29, respectively.

Given that *Pca* urediniospores are dikaryotic, the diploid aware assembler FALCON in combination with FALCON-Unzip (31) was used to first assemble the genomes of 12NC29 and 12SD80 and then distinguish regions of homology and divergence between haplotypes. Homologous regions were collapsed during FALCON assembly and are referred to as primary contigs, whereas divergent regions between haplotypes were assembled into haplotigs by FALCON-Unzip. As such, the primary contigs should contain the equivalent of one haploid genome and haplotigs represent the total sequence placed in alternate assembly paths relative to each individual primary contigs (**Figure 2A**). Genome assembly of 12SD80 resulted in 603 primary contigs with a total size of 99.2 Mbp and a contig N50 of 268.3 kbp, while 777 primary contigs with a total size of 105.2 Mbp and a contig N50 of 217.3 kbp were assembled for 12NC29 (**Table 1**). These assemblies demonstrate the advantage of long-read assembly to improve contiguity compared to previous short-read assemblies of other rust species. For example, the wheat stripe rust fungus, *Puccinia striiformis* f. sp. *tritici* (*Pst*), genome assembly contained more than 29,000 contigs with an N50 of 5.1 kbp (19) and the flax rust fungus, *Melampsora lini* (*Ml*), assembly has 21,000 scaffolds with an N50 of 31 kbp (22). The contiguity of our *Pca* genome assemblies are comparable to the scaffolding efficiency of the large insert Sanger sequence-based assemblies of the poplar rust fungus, *Melampsora larici-populina* (*Mlp*), and the wheat stem rust fungus, *Puccinia graminis* f. sp. *tritici* (*Pgt*), which contained 462 and 392 scaffolds, respectively (17). However, the *Mlp* and *Pgt* scaffolds contain approximately 3.5 and 7 Mbp of missing data, respectively, as gaps between contigs. The estimated genome sizes of 12SD80 and 12NC29 are in the range of other related rusts such as *Pgt* (92 Mbp) (17, 18) and *Pst* (65-130 Mbp) (19, 21, 24) and in agreement with nuclear DNA fluorescence intensity measurements of haploid pycniospores suggesting about 15% larger genome size of *Pca* relative to *Pgt* (32). Similarly, a preliminary genome assembly of another *Pca* isolate based on Illumina short-reads suggested a genome size of 110 Mbp (Park *et al*., unpublished). On the other hand, Tavares et al. (33) reported a haploid genome size of approximately 244 Mbp based on nuclear fluorescence for a *P. coronata* isolate obtained from *Avena sterilis*. Given the broad host range of *P. coronata (2)* this isolate may represent a different forma specialis.

**Figure 2.**
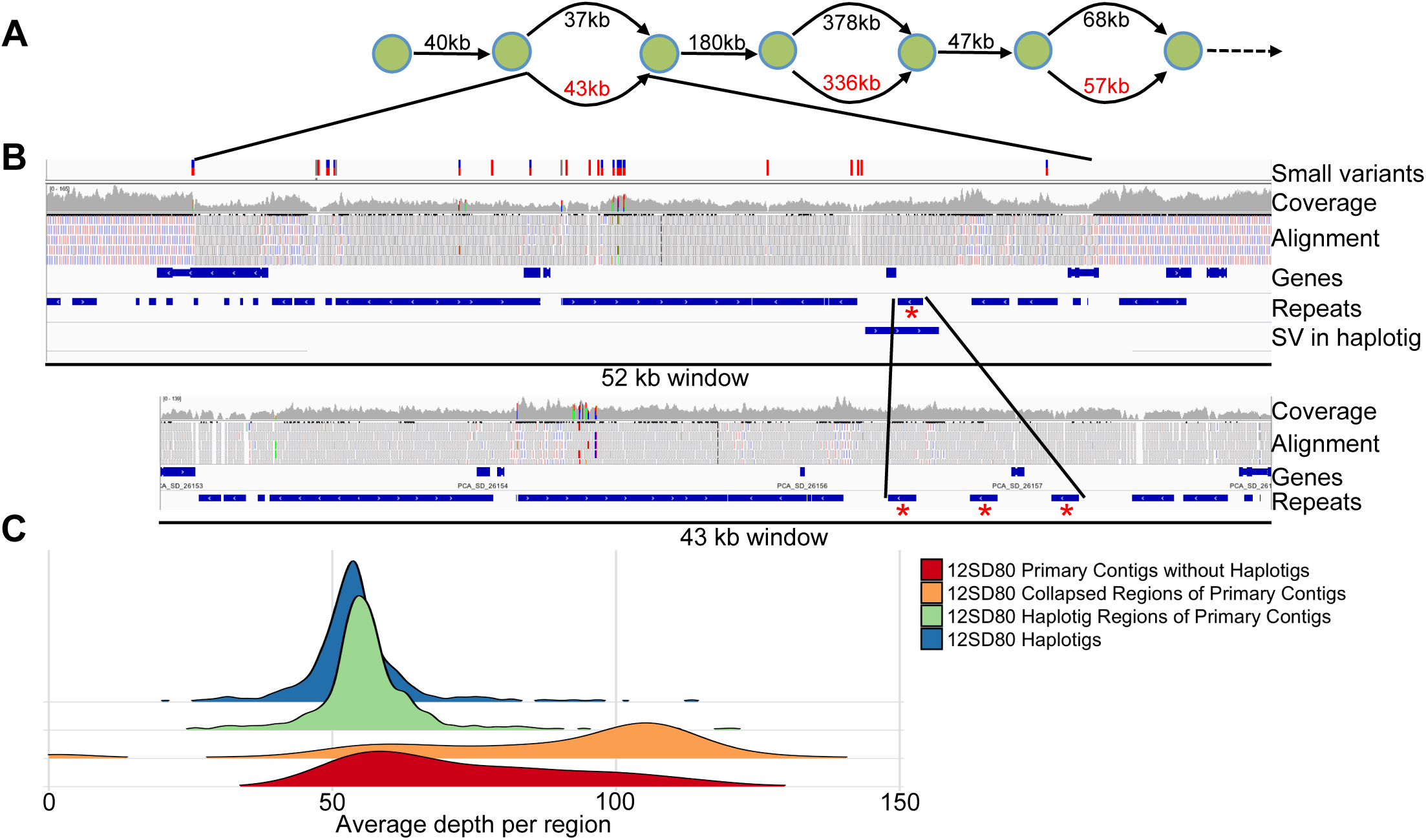
Characteristics of haplotig regions in a primary contig for the *Pca* isolate 12SD80. (**A**) Schematic depicting the first three haplotig regions of the largest primary contig in 12SD80 (000000F). The green circles represent nodes in the assembly graph and the numbers represent the distance between nodes for the primary contig (upper path, black) and haplotigs (lower path, red). (**B**) An IGV genome browser view of the first haplotig associated region of 12SD80 contig 000000F (upper panel) and the corresponding haplotig (lower panel). The top track shows SNPs and indels between haplotypes. The next track shows the coverage of short-read mapping to the assembly, and below that is the raw alignment evidence. Uniquely mapping reads are shown in red (-ve strand orientation) and blue (+ve strand) while grey indicates reads mapping to multiple locations. Annotated genes and repeats are shown in separate tracks, and the bottom track for the primary contig shows structural variations (SV). Red asterisks indicate a repeat element that has undergone a tandem expansion in the haplotig. (**C**) Density histograms of mean coverage depth of collapsed and haplotig regions of primary contigs, haplotigs, and primary contigs without haplotigs in 12SD80.

**Table 1.** Assembly metrics and evaluation

|  | <u>12SD80 Primary</u><br><u>Contigs</u> | <u>12SD80</u><br><u>Haplotigs</u> | <u>12NC29 Primary</u><br><u>Contigs</u> | <u>12NC29</u><br><u>Haplotigs</u> |
| --- | --- | --- | --- | --- |
| # contigs ($\geq 0$ bp) | 603 | 1033 | 777 | 950 |
| # contigs ($\geq 50000$ bp) | 475 | 372 | 560 | 403 |
| Total length (Mb) | 99.2 | 51.3 | 105.2 | 61.0 |
| Total length $\geq 50000$ bp (Mb) | 94.9 | 36.2 | 98.0 | 49.4 |
| Largest contig (Mb) | 1.39 | 0.35 | 1.19 | 0.48 |
| GC (%) | 44.7 | 44.9 | 44.7 | 44.9 |
| N50 (Kb) | 268.3 | 77.8 | 217.3 | 121.2 |
| Complete BUSCOs (%) | 90.4 | 57.9 | 89.6 | 72.1 |
| Complete and single-copy BUSCOs (%) | 85.9 | 57.2 | 84.1 | 69.7 |
| Complete and duplicated BUSCOs (%) | 4.5 | 0.7 | 5.5 | 2.4 |
| Fragmented BUSCOs (%) | 3.1 | 3.8 | 2.8 | 4.1 |
| Missing BUSCOs (%) | 6.5 | 38.3 | 7.6 | 23.8 |

A total of 1,033 and 950 haplotigs were assembled for 12SD80 and 12NC29, respectively, comprising 52% of the haploid genome size in each case (**Table 1**). Haplotig sequences were aligned to primary contigs to identify corresponding regions; illustrated for the largest primary contig in 12SD80 in **Figure 2A**. Numerous small variants were detected in the first haplotig-associated region in this primary contig and the corresponding haplotig by alignment of Illumina DNA reads to primary contigs and haplotigs simultaneously (**Figure 2B**). The haplotig also contains a tandem repeat expansion relative to the primary contig, while the flanking collapsed regions in the primary contig are less variable. The variation in this region likely explains why an alternate path in the assembly graph led to the phasing of this genomic region. The Illumina read depth (coverage) in the haplotig region is lower relative to the flanking collapsed regions as is expected considering that haplotig-associated regions represent a single haplotype, whereas most collapsed regions in primary contigs represent both haplotypes. In addition, reads in the collapsed region map uniquely in the genome, while those in the haplotig region map to multiple sites.

To validate haplotype phasing more extensively, we calculated genome-wide coverage for collapsed and haplotig-associated regions within primary contigs, as well as haplotigs. Haplotigs and haplotig regions of primary contigs in 12SD80 showed tight coverage distribution, with mean coverages of 56.3 and 58.7 respectively, while collapsed regions had a mean coverage of 103.6, but showed a broader distribution (**Figure 2C**). Regions of primary contigs with lower coverage but without an associated haplotig may represent locations with complex rearrangements or very large insertions/deletions between the two haplotypes. This could result in the presence of haplotype-specific sequences in primary contigs. Additionally, some primary contigs did not contain any associated haplotigs, which may be because the haplotype sequences were too divergent and assembled as two separate primary contigs. Consistent with this, primary contigs without haplotigs showed a lower coverage distribution than those with associated haplotigs (**Figure 2C**). Similarly, in 12NC29 mean coverages of haplotigs, haplotig regions and collapsed regions of primary contigs were 62.6, 64.3 and 91.0, respectively (**Figure S2A**). In 12SD80 and 12NC29, there were 176 and 312 primary contigs without haplotigs, respectively, totaling 11.1 and 17.5 Mbp. If these do represent separately assembled haplotypes, then this may partly explain the approximately 6 Mbp larger primary contig assembly size for 12NC29. The ability to phase the genome assembly into primary contigs and haplotigs in this fashion represents a significant advance to compare haplotype composition in dikaryotic fungi.

### Assessment of genome completeness and repetitive DNA content

To assess the completeness of the *Pca* genome assemblies, highly conserved fungal genes were mapped in the primary contigs and haplotigs using BUSCO (34). Approximately 90% of the BUSCO genes were present as complete sequences and nearly an additional 3% as fragmented copies in the primary contigs of both genome assemblies (Table 1). One additional BUSCO gene not present in the primary contigs was found on a haplotig in 12SD80, while no unique BUSCO genes were found in 12NC29 haplotigs. Fourteen BUSCOs (4.8%) were missing in both isolates, which suggests the presence of difficult to assemble regions in the *Pca* genome. A search for telomere repeat sequence at the ends of all contigs detected 11 unique telomeres in 12NC29 and 15 in 12SD80, out of an estimated 16–20 chromosomes (35). Overall, these results indicated that the primary contigs are a good representation of the core dikaryotic genome of *Pca*.

RepeatMasker detected interspersed repeats covering about 53% of the assembled *Pca* genomes (primary contigs and haplotigs combined; **Table 2**), similar to other rust fungi which are typically in the range of 35-50% (17, 21, 22). The most prevalent repetitive elements belonged to the LTR retroelement class (20% of the genome), which was also found to be the most abundant class in *Pgt* and *Mlp* (17, 24), while DNA elements accounted for about 15% of the genome. The GC content was approximately 45% for primary contigs and haplotigs in both *Pca* isolates (**Table 1**), which is consistent with other rust species, such as *Ml* (41%) (22). The distribution of GC content in individual contigs (**Figure S2B**) did not display a bimodal distribution which would indicate the presence of AT-rich regions, such as those observed in fungi that use repeat-induced point mutation (RIP) to inhibit transposon proliferation (36).

**Table 2.**
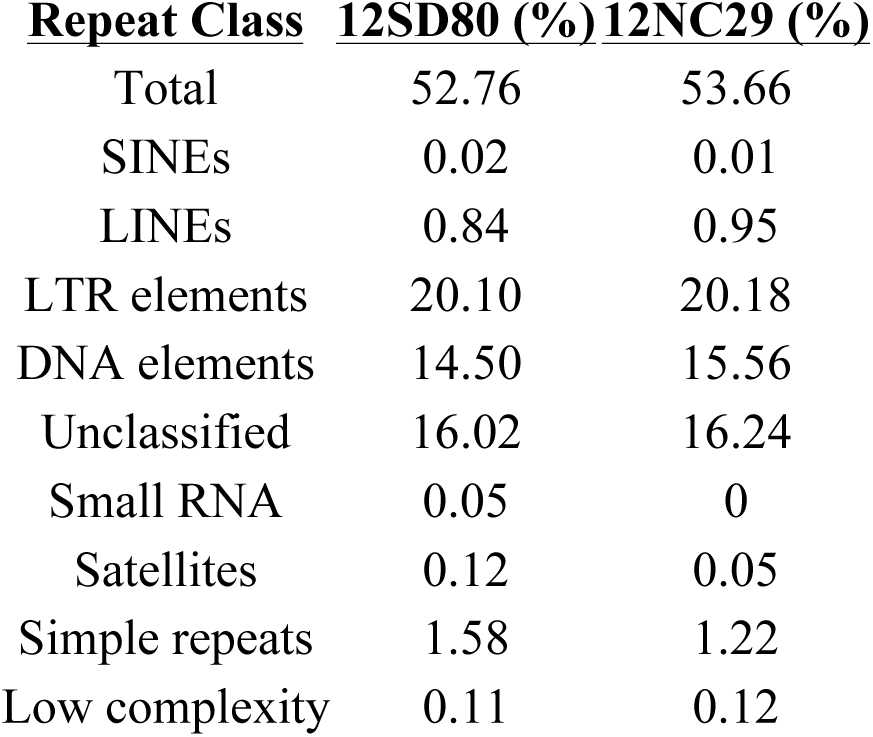
Proportion of repeated sequence content in *Pca* isolates

| **<u>Repeat Class</u>** | **<u>12SD80 (%)</u>** | **<u>12NC29 (%)</u>** |
| --- | --- | --- |
| Total | 52.76 | 53.66 |
| SINEs | 0.02 | 0.01 |
| LINEs | 0.84 | 0.95 |
| LTR elements | 20.10 | 20.18 |
| DNA elements | 14.50 | 15.56 |
| Unclassified | 16.02 | 16.24 |
| Small RNA | 0.05 | 0 |
| Satellites | 0.12 | 0.05 |
| Simple repeats | 1.58 | 1.22 |
| Low complexity | 0.11 | 0.12 |

### Gene annotation and orthology prediction revealed phased allele pairs within isolates and orthologs between isolates

For each *Pca* isolate, RNAseq reads from germinated spores, isolated haustoria and infected oat leaves at 2 and 5 dpi (**Table S2**) were pooled and used to generate both *de novo* and genome-guided transcriptome assemblies using Trinity v2.4.0 (37). These assemblies were used as transcriptional evidence in the Funannotate pipeline along with alignment evidence from publicly available EST clusters for Pucciniamycotina species. In total, 17,248 and 17,865 genes were annotated on primary contigs for 12SD80 and 12NC29, respectively (**Table 3**), which is similar to the haploid gene content of other rust fungal genomes (17, 22). An additional 9,548 and 10,936 genes were annotated on haplotigs for 12SD80 and 12NC29, respectively.

**Table 3.** Gene, allele and ortholog content in *Pca* genome assemblies

|  | <u>12SD80</u> | <u>12NC29</u> |
| --- | --- | --- |
| Total genes (P and H*) | 26,796 | 28,801 |
| Mean gene length (all genes) | 1,516 bp | 1,518 bp |
| % of genome covered by genes | 27.0 | 26.3 |
| Total genes on P | 17,248 | 17,865 |
| Total genes on H | 9,548 | 10,936 |
| Allele pairs on P and H | 6,664 | 7,789 |
| Haplotype-singleton genes on P | 2,162 | 2,311 |
| Haplotype-singleton genes on H | 2,033 | 2,154 |
| Effectors on P | 529 | 549 |
| Effectors on H | 320 | 351 |
| Effectors on P in allele pairs | 268 | 277 |
| Effectors on H in allele pairs | 262 | 276 |
| Haplotype-singleton effectors on P | 42 | 61 |
| Haplotype-singleton effectors on H | 49 | 62 |
| Orthologous effectors on P between isolates | 336 | 327 |
| Isolate singleton effectors on P | 184 | 216 |

\* P and H represent primary contigs and haplotigs, respectively.

To identify putative allele pairs in the phased assemblies, we searched for genes on primary contigs that had an ortholog present on the corresponding haplotig using Proteinortho (38) in synteny mode to account for gene order (**Table 3**). A total of 6,664 and 7,789 such allele pairs were identified in 12SD80 and 12NC29, respectively. About 2,000 haplotype-singletons, with no orthologs in a corresponding region, were also detected in haplotig-regions of primary contigs, with a similar number in haplotigs (**Table 3**). These singletons represent haplotype-specific genes with presence/absence variation or genes with substantial sequence variation that prevents orthology detection. We also examined gene orthology between isolates, and identified 9,764 orthologous groups (∼55% of all genes) containing either: 1) two orthologous genes, one from each isolate with no allele pairs, 2) an allele pair from one isolate with an unpaired gene from the other, or 3) two allele pairs, one from each isolate. Isolate-singletons may represent presence/absence polymorphisms or could be due to sequence divergence or genome rearrangements preventing orthology detection. Therefore, we examined gene coverage by cross-mapping Illumina reads from each isolate onto the other assembly (**Figure S3**). The isolate-singleton genes in 12SD80 and 12NC29 included 558 and 1,174 genes, respectively, with low coverage (<30X) suggesting they represent presence/absence polymorphisms, while the remainder showed higher coverage (30 – 200X) indicating that homologs may be present in both isolates. Taken together, these findings indicate a high level of gene content variation between haplotypes and isolates of *Pca*. Sequencing a larger sample of *Pca* isolates will help determine the number of conserved (core) genes versus isolate-specific genes in this species.

### Functional annotation of *Pca* genomes

GO term abundances of annotated genes on primary contigs and haplotigs combined were very similar between isolates with no significant GO term enrichments or depletions. Examination of KEGG pathway annotations (39) indicated that, as observed for other rust fungi (17, 22, 24), the *Pca* genomes lacked nitrate and nitrite assimilation genes. The assemblies did contain the enzymes glutamine synthetase (K01915), glutamate synthase (K00264), and glutamate dehydrogenase (K00260), which are putatively involved in nitrogen assimilation from host-derived amino acids. Enzymes of the sulfate assimilation pathway were also absent in the two *Pca* isolates. Notably, sulfite reductase was missing from both assemblies, as was observed for *Pgt* (17). These observations are consistent with the loss of nitrate, nitrite, and sulfate assimilation pathways associated with the evolution of obligate biotrophy in rust fungi (17, 22). Most categories of transcription factor families showed low abundance in both isolates except the CCHC zinc finger class (IPR001878) that has 103 members in 12NC29 and 48 in 12SD80 (**Figure 3A**). This family was also expanded in *Pgt* and *Mlp* relative to other fungi (17) and are of particular interest as zinc finger TFs are hypothesized to play roles in effector regulation (40).

**Figure 3.**
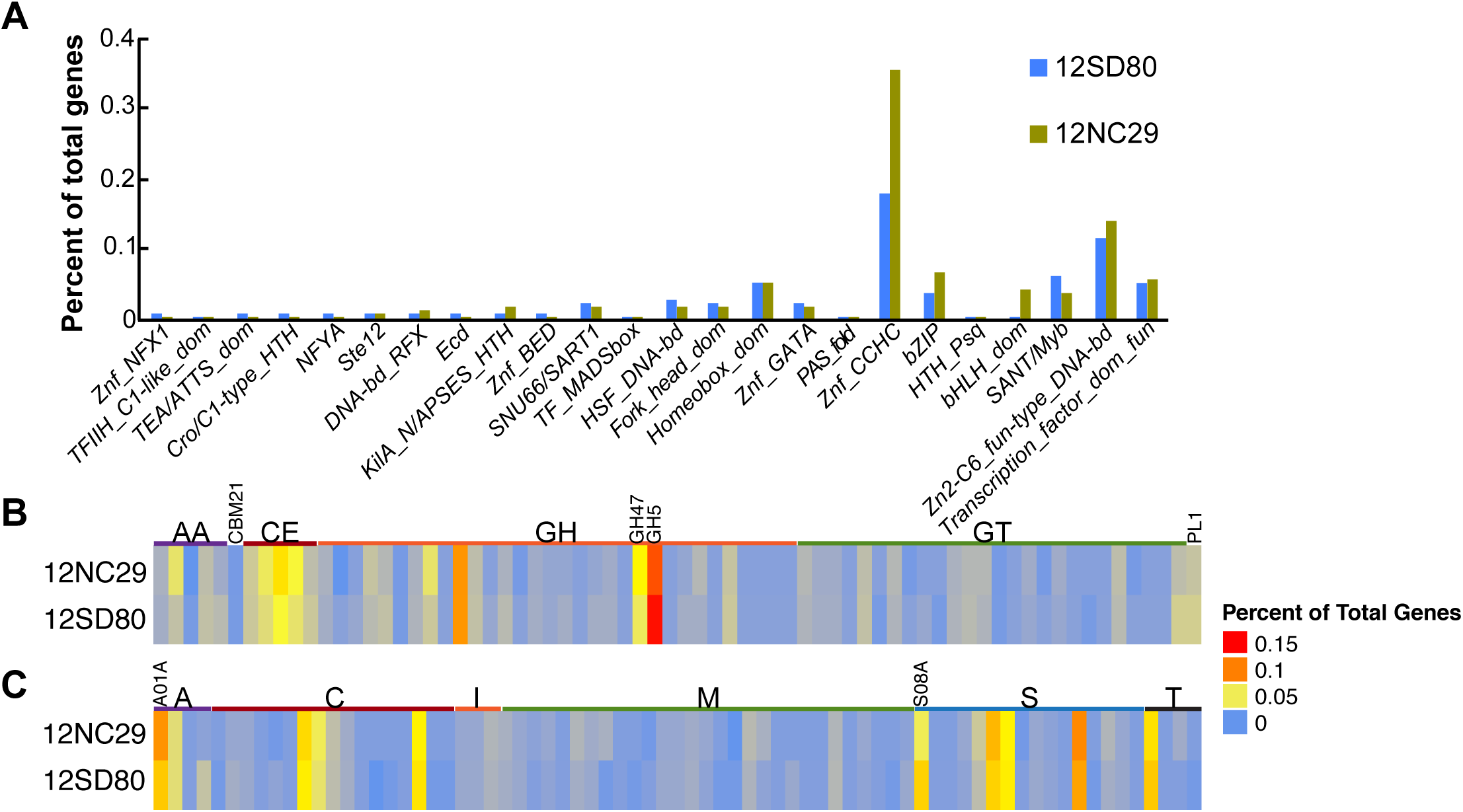
Functional annotation of transcription factors, CAZymes, and MEROPS proteases in *Pca* isolates. (**A**) Percent of total genes predicted to encode members of various fungal transcription factor classes based on InterProScan annotation. (**B**) Heatmap showing percent of total genes annotated as members of CAZyme families in the following classes: auxiliary activities (AA), carbohydrate-binding modules (CBM), carbohydrate esterases (CE), glycoside hydrolases (GH), glycosyltransferases (GTs), and polysaccharide lyases (PL). Expanded families GH5 and GH47 are indicated. **C**) Heatmap showing percent of total genes annotated as members of MEROPS families of aspartic acid (A), cysteine (C), metallo (M), serine (S), and threonine (T) proteases or peptidase inhibitors (I). Expanded families A01A and S08A are indicated.

### Heterozygosity in the dikaryotic genome of *Pca*

Heterozygous small variants, including single-nucleotide polymorphisms (SNPs), insertions/deletions (indels) and multiple-nucleotide polymorphisms (MNPs), were identified by mapping Illumina reads to only primary contigs in each isolate. We detected 3.45 and 4.60 heterozygous variants/kbp (including 2.68 and 3.62 SNPs/kbp) in 12SD80 and 12NC29, respectively. These heterozygosity rates are in line with genome-wide estimates of 1-15 hetSNPs/kbp for other *Puccinia* spp. (18, 19, 21, 24), although such estimates may be influenced by differences in variant calling methods and parameters, residual assembly errors, read length and coverage, and may differ between isolates of a species. When Illumina reads from 12SD80 were mapped to the 12NC29 primary contig reference, we detected a total of 3.48 heterozygous and 2.31 homozygous variants/kbp. In the reciprocal comparison, 5.60 heterozygous and 1.75 homozygous variants/kbp were identified, indicating substantial variation between isolates as well as between haplotypes.

The majority of variants between haplotypes were found in intergenic regions (**Figure S4A**), and these occurred at a higher frequency (3.66 and 4.88 variants/kbp in 12SD80 and 12NC29, respectively) than variants in genic regions (2.86 and 3.76 variants/genic kbp). Heterozygosity rates were higher in haplotig regions of primary contigs (4.36 and 5.50 variants/kbp in 12SD80 and 12NC29, respectively) than collapsed regions (1.06 and 1.27 variants/kbp). These observations are consistent with haplotigs containing regions of divergence between haplotypes.

We also compared heterozygosity rates in *Pca* and the rust species *Mlp*, *Ml*, *Pst*, and *Pt* using a *k-mer* profile approach based on available Illumina reads with the software GenomeScope (41). In this analysis, homozygous genomes display a simple Poisson distribution in the *k-mer* profile plots, whereas heterozygous genomes give a bimodal profile. The *k-mer* profiles of most of these species (**Figure S5**) showed bimodal profiles, which indicated fairly heterozygous genomes. This was less apparent for *Pst* and *Ml*, which may be explained by the shorter-read lengths and lower coverage datasets for these species. Heterozygosity levels calculated in this analysis were similar for all species, but lower than levels detected by SNP calling.

To assess structural variation (SV) between haplotypes we compared haplotigs to their corresponding aligned regions in primary contigs using Assemblytics, which detects three types of SV: large insertions/deletions; tandem expansions/contractions, which involve tandemly repeated sequences; and repeat expansions/contractions in which homologous regions are separated by regions with no homology in each sequence (42). The distributions of these classes of SV are very similar between the two isolates (**Figure S6**), with insertions/deletions and repeat expansions/contractions more prevalent than tandem expansions/contractions. Such SV between 50 and 10,000 bp in size represented 2.7% of the primary contig genome size in 12NC29 and 2.1% in 12SD80, and impacted 646 and 951 coding regions on primary contigs in 12SD80 and 12NC29, respectively (**Figure S4B**).

### Prediction of secretome and candidate effectors

Pathogen effectors are secreted proteins that manipulate host cell processes to facilitate infection, but can also be recognized by host resistance genes (43). Thus, differences in virulence profiles between 12NC29 and 12SD80 (**Figure 1A**) likely result from variation in their effector repertoires. We predicted 1,532 and 1,548 secreted proteins on primary contigs of 12SD80 and 12NC29, respectively, corresponding to about 9% of all protein-coding genes. Similarly, 941 and 1,043 secreted proteins were predicted on haplotigs in 12SD80 and 12NC29, respectively, (including 773 and 856 in allele pairs). About 35% of all secreted proteins were predicted as effectors by the EffectorP machine learning tool for fungal effector prediction (44) (**Table 4**). No enriched GO terms were detected among the predicted effectors, and the vast majority had no homologs with known or predicted function (**Table S3**), as is commonly observed for fungal effectors (45).

**Table 4.** Features of proteins encoded by genes in different expression clusters of *Pca*

| Clusters | # proteins | %CAZymes | %EffectorP | %NLS (LOCALIZER) | %ApoplastP |
| --- | --- | --- | --- | --- | --- |
| <u>12SD80</u> |  |  |  |  |  |
| 1 | 251 | 18.7 | 37.1 | 20.7 | 23.1 |
| 2 | 78 | 6.7 | 29.5 | 19.2 | 43.6 |
| 3 | 55 | 6.7 | 27.3 | 12.7 | 61.8 |
| **4\*** | **111** | **2.7** | **35.1** | **30.6** | **8.1** |
| **5\*** | **173** | **5.3** | **36.4** | **24.9** | **16.8** |
| 6 | 197 | 36.0 | 20.3 | 18.8 | 30.5 |
| 7 | 198 | 24.0 | 42.9 | 16.7 | 48.0 |
| <u>12NC29</u> |  |  |  |  |  |
| 1 | 239 | 17.1 | 36.0 | 23.8 | 25.1 |
| 2 | 93 | 14.5 | 33.3 | 30.1 | 30.1 |
| **3\*** | **129** | **6.6** | **41.9** | **19.4** | **10.1** |
| 4 | 71 | 7.9 | 23.9 | 9.9 | 53.5 |
| 5 | 166 | 19.7 | 24.1 | 22.9 | 28.3 |
| **6\*** | **179** | **5.3** | **36.3** | **27.9** | **20.1** |
| 7 | 86 | 7.9 | 45.3 | 10.5 | 52.3 |
| 8 | 124 | 13.2 | 29.0 | 18.5 | 37.1 |
| 9 | 60 | 7.9 | 26.7 | 8.3 | 55.0 |

\* Red indicates haustorially-expressed clusters.

RNAseq datasets from different tissue types were used to identify secreted protein genes in primary contigs of each isolate that were differentially expressed during infection, and similarly expressed genes were grouped using *k*-means clustering. This analysis detected seven distinct expression profile clusters for 12SD80 and nine for 12NC29 (**Figure 4A** and **B**, **Table 4**). Genes in clusters 4 and 5 in 12SD80 showed high expression in haustorial samples and also relatively high expression in infected leaves, with those in cluster 4 showing the lowest expression in germinated urediniospores. Similar profiles were observed for clusters 3 and 6 in 12NC29. These expression patterns are consistent with those of previously identified secreted rust effectors that enter host cells, which show high expression in haustoria (5). About 35-40% of the secreted genes in these clusters were predicted as effectors by EffectorP (**Table 4**). These clusters also show relatively high proportions of genes encoding predicted nuclear localized proteins and the lowest proportions of apoplast localized proteins as predicted by ApoplastP (Sperschneider *et al*., submitted for publication) (**Table 4**), suggesting that these clusters are enriched for host-delivered effectors.

**Figure 4.**
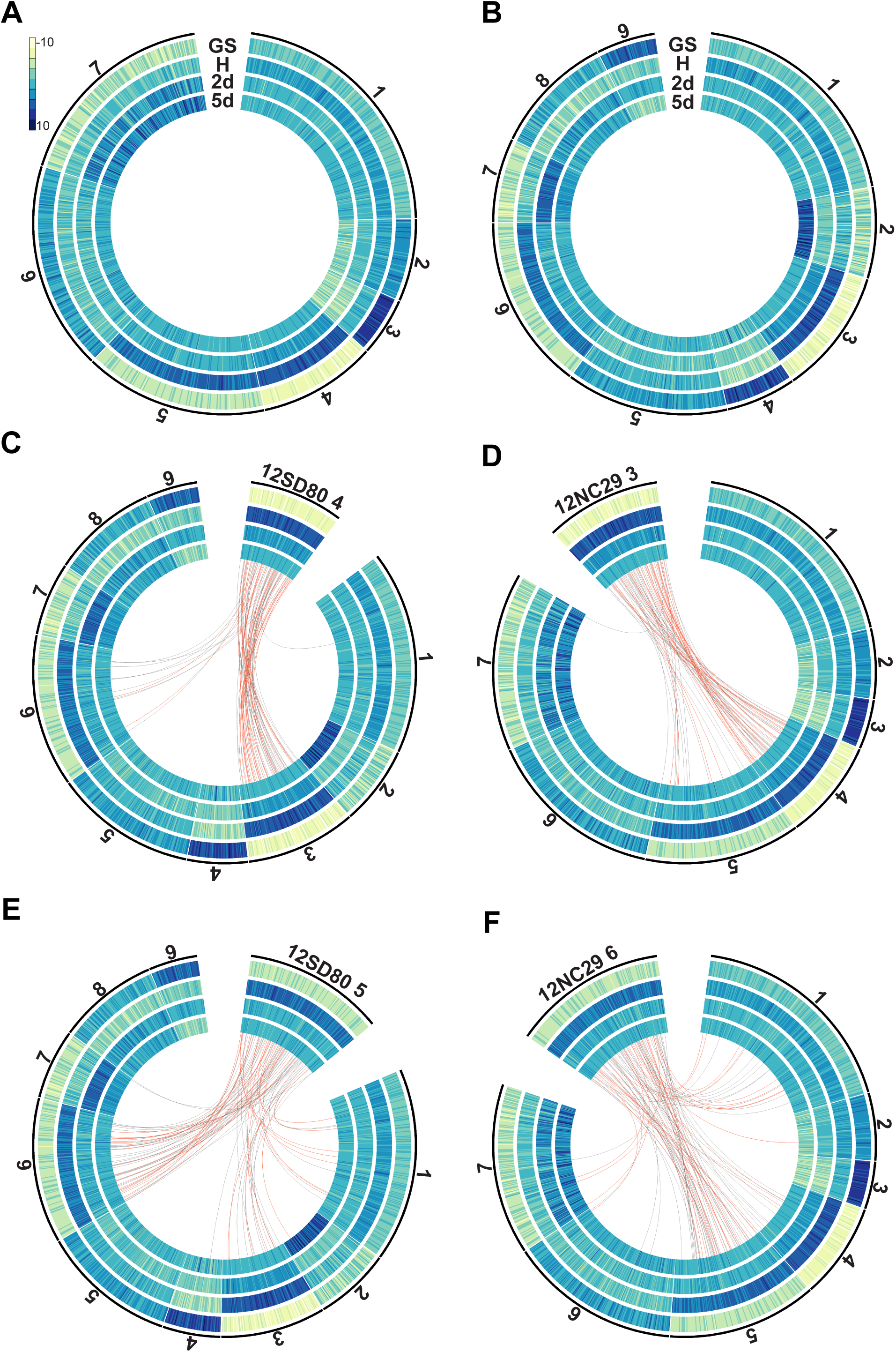
Clustering analysis of predicted secretome gene expression profiles and orthology in *Pca*. (**A**) *K-*means clustering of secretome genes of 12SD80 and (**B**) 12NC29. Heatmaps show rlog transformed expression values with dark blue indicating high expression according to the scale. Cluster numbers are shown outside of the graphs and tracks show gene expression in germinated spores (GS), isolated haustoria (H), and infected tissues at 2 (2d) and 5 dpi (5d). (**C**) Orthology relationships between genes in 12SD80 cluster 4 and all 12NC29 clusters are indicated by red (predicted effectors) and grey (other secreted proteins) lines. (**D**-**F**) Orthology relationships between genes in 12NC29 cluster 3 and all 12SD80 clusters (**D**), 12SD80 cluster 5 and all 12NC29 clusters (**E**), and 12NC29 cluster 6 and all 12SD80 clusters (**F**).

GO analysis detected an enrichment for molecular functions related to glycosyl hydrolase and peptidase activities in the *Pca* secretome (**Figure S7**), which may indicate roles for these proteins during infection in the plant apoplast. Necrotrophic and hemibiotrophic plant pathogenic fungi secrete large numbers of carbohydrate-active enzymes (CAZymes) including plant cell wall-degrading enzymes (PCWDEs) that are important for host invasion (46-48). However, biotrophs such as rust fungi contain far fewer of these enzymes and their roles are less well defined, although roles in both plant cell wall degradation and fungal cell wall reorganization have been suggested based on expression data for *Mlp* and *Pgt (49)*. We detected 350 and 374 CAZymes in isolates 12SD80 and 12NC29, respectively, of which about 20% (75 and 76 CAZymes) were predicted to be secreted. This is consistent with estimates for other biotrophs from a fungal kingdom-wide analysis of secreted proteins (50). Secreted CAZymes were most abundant in expression cluster 6 in 12SD80 (36%) and cluster 5 in 12NC29 (20%), which both showed slightly elevated expression in germinated spores, but also significant expression under *in planta* conditions (**Table 4, Figure 4A** and **B**), suggesting that these enzymes have roles throughout development. Interestingly, the clusters with the strongest expression in germinated spores compared to other conditions (cluster 3 in 12SD80, and clusters 4 and 9 in 12NC29) have relatively low proportions of CAZymes and the highest percentage of predicted apoplast-localized proteins. This may indicate that *Pca* employs a repertoire of apoplastic effectors that do not have similar enzymatic function to CAZymes.

Glycoside hydrolase (GH) enzymes are a subclass of CAZymes, with 175 and 182 members detected in 12SD80 and 12NC29, respectively (**Figure 3B**). Of these, 43 and 46 were predicted to be secreted in 12SD80 and 12NC29, respectively representing approximately 60% of all secreted CAZymes. The GH5 (cellulase and other diverse enzymatic functions are in this family) and GH47 (α-mannosidases) families were expanded in *Pca*, as seen in *Pgt* and *Mlp* (17), with 32 GH5 family members in both isolates, and 13 and 18 GH47 family members in 12SD80 and 12NC29, respectively. However, only 2-4 members of these families were predicted as secreted, suggesting that these families have mostly intracellular roles. Consistent with previous observations in rust fungi (17) the cellulose-binding module 1 subfamily (CBM1) was not found in *Pca.*

Secreted subtilases (serine proteases) and aspartic proteases are predicted to act as effectors in rust fungi and may interfere with plant defense responses (51, 52). Both the A01A (aspartic proteases) and S08A (subtilisin-like serine proteases) families were expanded in the *Pca* genomes as was found for *Pgt* and *Mlp* (17) (26 and 34 members of A01A and 25 and 18 members of S08A in 12SD80 and 12NC29, respectively, **Figure 3C**). A total of 11 (42%) and 17 (50%) aspartic proteases and 17 (68%) and 15 (83%) serine proteases are predicted to be secreted in 12SD80 and 12NC29, respectively. Unlike secreted CAZymes, these secreted proteases have no obvious clustering pattern amongst differentially expressed secretome genes.

**Variation in effector candidates**. Similar to genome-wide patterns, heterozygous small variants were more abundant in 1,000 bp upstream and downstream regions than transcribed regions of effector candidate genes (**Figure S4C**). The rate of heterozygous variants was slightly higher in effectors on primary contigs compared to all genes on primary contigs in 12NC29, but not in 12SD80, as was the nonsynonymous variant rate (**Table 5**). Elevated variation rates in effector genes relative to all genes were also observed in between isolate comparisons. SV impacted 13 and 23 predicted effectors on primary contigs in 12SD80 and 12NC29, respectively (**Figure S4D**) including examples of presence/absence and copy number variation.

**Table 5.**
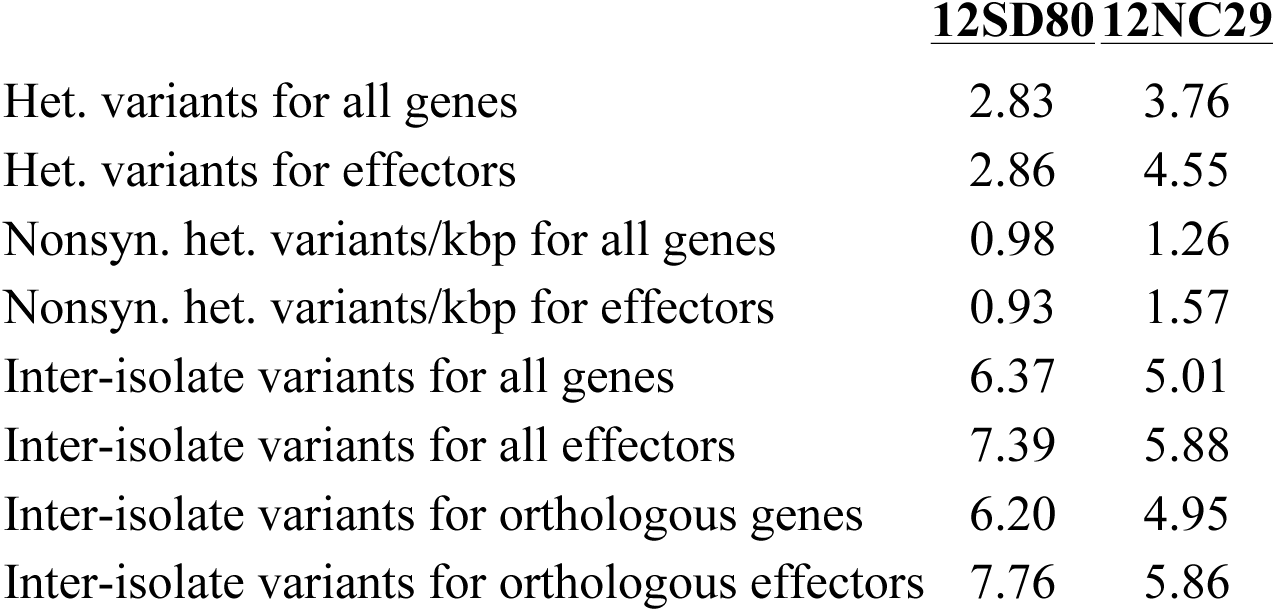
Variation rates (variants/kbp) in annotated genes and predicted effectors on primary contigs in *Pca*.

|  | <u>12SD80</u> | <u>12NC29</u> |
| --- | --- | --- |
| Het. variants for all genes | 2.83 | 3.76 |
| Het. variants for effectors | 2.86 | 4.55 |
| Nonsyn. het. variants/kbp for all genes | 0.98 | 1.26 |
| Nonsyn. het. variants/kbp for effectors | 0.93 | 1.57 |
| Inter-isolate variants for all genes | 6.37 | 5.01 |
| Inter-isolate variants for all effectors | 7.39 | 5.88 |
| Inter-isolate variants for orthologous genes | 6.20 | 4.95 |
| Inter-isolate variants for orthologous effectors | 7.76 | 5.86 |

Orthologous gene relationships for effectors were identified to examine the conservation of effector repertoires between haplotypes and isolates. Approximately 50% of predicted effectors had an allele pair (**Table 3, Dataset S1** to **S4**), while a total of 91 (11%) and 123 (14%) predicted effectors were haplotype singletons in 12SD80 and 12NC29, respectively (**Table 3, Dataset S5** to **S8**). For 12SD80, 336 predicted effector genes on primary contigs had orthologs in 12NC29 (primary contigs and haplotigs), while 184 were isolate-singletons, with similar numbers observed for the reciprocal comparison (**Table 3, Dataset S9** - **S12**). Inter-isolate variation rates in orthologous effector genes were slightly elevated when compared to all orthologous genes (**Table 5**). Overall, these results showed substantial variation in effector gene candidates both between haplotypes and isolates that may provide a basis for virulence differences between the isolates.

### Conservation of expression patterns between orthologous secreted proteins

When orthology relationships were overlaid onto the secretome expression clusters for each isolate, the majority of orthologous secreted proteins and predicted effectors showed conserved expression patterns between 12SD80 and 12NC29 (**Figure 4C-F, Figures S8** and **S9**). For instance, orthologs of genes in cluster 4 of 12SD80 with the strongest haustorial expression relative to germinated spores were mainly found in cluster 3 in 12NC29, which showed an equivalent expression profile (**Figure 4C**). A number of orthologs were also found in 12NC29 cluster 6, which shows the next strongest haustorial expression, while there was a single ortholog in 12NC29 cluster 1, which was slightly upregulated in haustoria compared to all other conditions. Similar conservation of expression profiles were observed for 12NC29 genes in cluster 3, which showed strong conservation of expression patterns to 12SD80 clusters 4 and 5 (**Figure 4D**). Genes in 12SD80 cluster 5 (the second strongest haustorial cluster) mostly showed orthology to genes in the equivalent cluster 6 in 12NC29, although some orthologs were in clusters 1 and 3 (**Figure 4E**). For 12NC29 cluster 6, a similar trend of expression conservation to 12SD80 cluster 5 was observed (**Figure 4F**). A few orthologous effector candidates showed divergent expression patterns between isolates. For instance, one effector in 12SD80 cluster 5 had an ortholog in 12NC29 cluster 4, which has the highest expression in germinated spores and another had an ortholog in cluster 2 showing highest expression at 5 dpi (**Figure 4E**). Such expression differences may contribute to differences in virulence phenotypes. Thus, future investigation of differential expression of orthologous effectors, as well as isolate-singleton effectors, may provide key insights into the mechanisms for virulence in *Pca*.

### Genomic context of predicted effector candidate genes

Genome sequences of several filamentous plant pathogens have provided evidence for a ‘two-speed genome’ model, in which rapidly evolving effector genes are preferentially located in low gene density and repeat rich regions (53). This genome architecture may favor fast host adaptation by relieving constraints on effector diversification. To determine the distribution of genes in gene-rich or sparse regions, we used a two-dimensional genome-binning method (54) to plot intergenic distances for all genes in *Pca* (**Figure 5**). Predicted effectors on primary contigs and haplotigs in both isolates showed no difference in location compared to the overall gene space. Moreover, both orthologous effector genes and isolate-singletons had similar intergenic distances to all genes. Genome-wide geometric correlation with the GenometriCorr R package (55) found no significant association between effector genes and repeat elements in either isolate. Thus, these findings do not support the presence of a ‘two speed genome’ in *Pca*, consistent with observations for other rust fungi (56).

**Figure 5.**
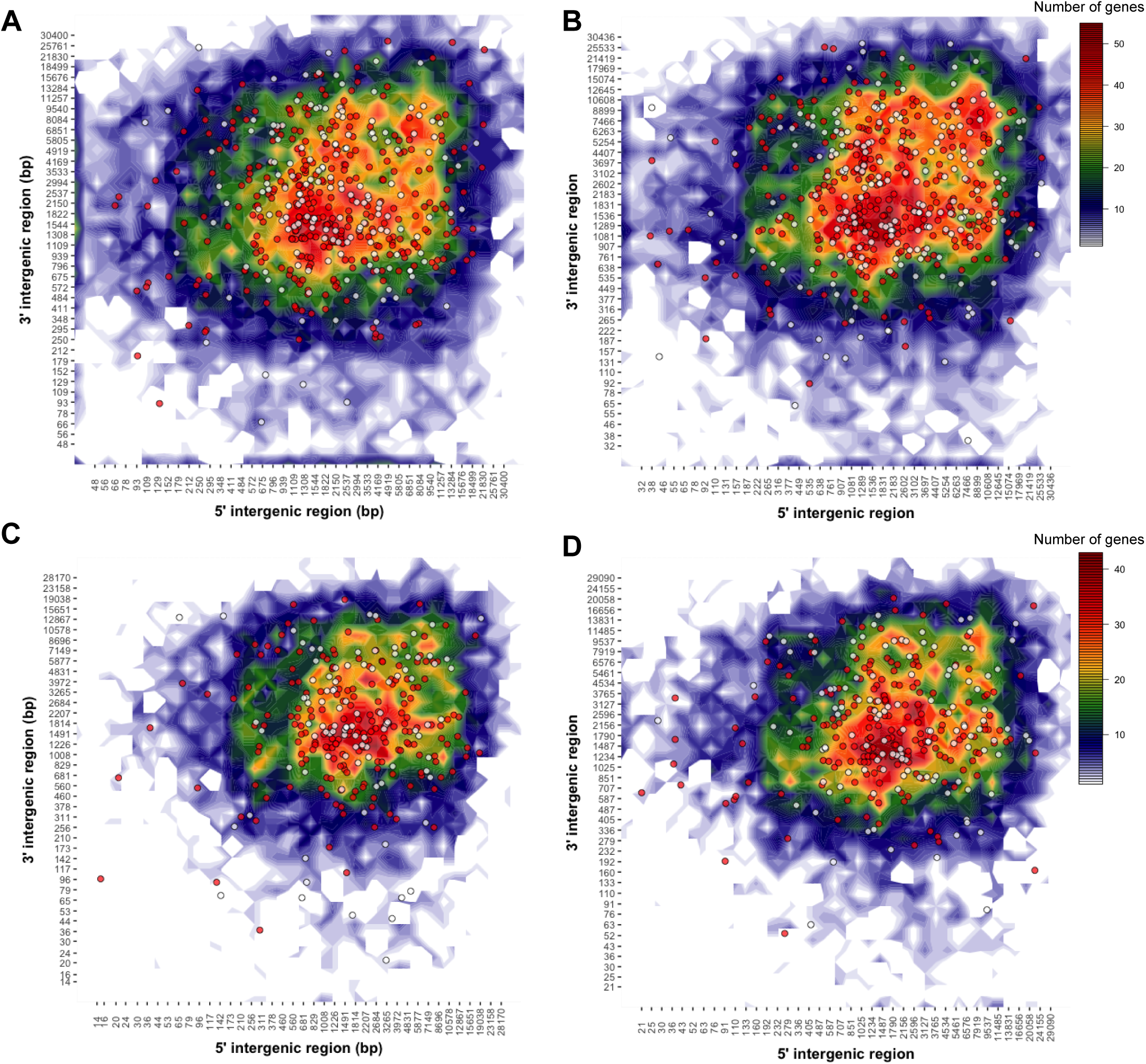
Genomic landscape of predicted *Pca* effectors. Heatmap plots representing the distribution of 5’ and 3’ intergenic distances for all genes on primary contigs of (**A**) 12SD80 and (**B**) 12NC29, and haplotigs of (**C**) 12SD80 and (**D**) 12NC29. Scales representing gene content per bin are shown on the right. Circles indicate predicted effectors with orthologs (red) or isolate-singletons (white).

## Conclusions and future directions

A significant challenge when assembling dikaryotic fungal genomes is to capture and align haplotype variation. Here, we demonstrate successful implementation of the diploid-aware long-read assembler FALCON and FALCON-Unzip to generate highly contiguous genome assemblies and resolve haplotypes from SMRT sequencing data for the oat crown rust fungus, *Pca*. These phased-assemblies allowed detection of structural variation between haplotypes equivalent to more than 2% of the genome size that impacted a significant number of genes and predicted effectors. This type of variation has not been previously examined in rust species due to the limitations imposed by collapsed short-read genome assemblies. Furthermore, the long-read assembly approach greatly improved contiguity compared to short-read assemblies of other rust fungi, which are highly fragmented due to an abundance of repetitive sequences in their genomes. Orthology analysis also allowed detection of allele pairs on the different haplotypes, as well as many genes potentially unique to one haplotype or highly diverged. We also observed high divergence in gene content and sequence between isolates, which may reflect their origins from geographically separated populations (South Dakota vs North Carolina). Transcriptome profiling revealed clusters of haustorially-expressed secreted proteins that are likely enriched for host-delivered effectors, as well as clusters of predicted CAZymes and apoplastic effectors that are preferentially expressed in germinated urediniospores.

Several mechanisms including mutation, sexual recombination and somatic hybridization are postulated to cause changes in virulence phenotypes in rust fungal populations (14, 16). However, few studies have specifically characterized molecular events associated with virulence variation, and large-scale whole-genome comparative population analyses have not been conducted for rust fungi. The high quality haplotype-phased genome references for two dikaryotic *Pca* isolates developed in this study provide the foundation for large-scale resequencing of *Pca* isolates to identify genetic variation underlying variability in virulence phenotypes. The identification of the *Avr* genes corresponding to known oat *R* genes will help to prioritize and pyramid broadly effective *R* genes in oat breeding programs.

## Materials and Methods

### *Puccinia coronata* f. sp. *avenae* (*Pca*) isolates and plant inoculations

*Pca* isolates 12NC29 (pathotype LBBB) and 12SD80 (pathotype STTG) were collected from North Carolina and South Dakota, respectively, by the USDA-ARS Cereal Disease Laboratory (CDL) annual rust surveys in 2012 and stored at −80°C. To ensure isolate purity, two single-pustule purifications from low density infections on seven-day old oat seedlings (variety ‘Marvelous’) were completed prior to amplification of urediniospores as described by Carson (6). Heat shock activated (45°C, 15 minutes) urediniospores were resuspended in Isopar M oil (ExxonMobil) at 2 mg spores/ml and for spray-inoculation (50 μl per plant). Inoculated plants were placed in dew chambers in the dark overnight (16 hours) with 2 minutes of misting every 30 minutes then maintained in isolated growth chambers (18/6 hour light/dark, 22/18°C day/night, 50% relative humidity). Pathotype assignment and final assessments of identity and purity of each isolate was performed using standard oat differential lines (2, 7), with infection scores converted to a 0-9 numeric scale for heat map generation.

### DNA extraction from *Pca* urediniospores for Illumina and PacBio Sequencing

Freshly harvested urediniospores were germinated as described (57) and fungal mats were vacuum dried, lyophilized and stored at −80ºC. The lyophilized tissue was ground in liquid nitrogen in 20-30 mg batches in 2 ml microcentrifuge tubes. DNA was extracted using genomic-tip 20/G columns (Qiagen catalog number 10223) following a user-supplied protocol (<u>https://www.qiagen.com/us/resources/resourcedetail?id=cb2ac658-8d66-43f0-968e-7bb0ea2c402a&lang=en</u>) except that lysis buffer contained 0.5 mg/ml of lysing enzymes from *Trichoderma harzianum* (Sigma L1412) and DNA was resuspended in Qiagen EB. Qubit (Invitrogen) and pulsed-field gel electrophoresis with a CHEF-DR III (Bio-Rad) were used to evaluate DNA quantity and quality, with yields of 15-20 ug per 200 mg of tissue obtained.

### Genomic DNA sequencing and *de novo* assembly

Approximately 10 μg of genomic DNA was purified with AMPure XP beads (Beckman Coulter) and sheared to an average size of 20 kbp using g-TUBEs (Covaris). Size and quantity were assessed using the TapeStation 2200 (Agilent Technologies). Library preparation followed the PacBio standard 20 kbp protocol, with size selection performed using a BluePippin (Sage Science) with a 0.75% agarose cassette and a lower cutoff of 7 kbp. Twenty five SMRT cells per library were run on the PacBio RSII (Pacific Biosciences) using P6/C4 chemistry, 0.15 nM MagBead loading concentration, and 360-minute movie lengths at the Frederick National Laboratory for Cancer Research (Frederick, MD, USA). Illumina libraries were prepared from 100 ng of genomic DNA with the TruSeq Nano DNA procedure and a 350 bp insert size. Both libraries were multiplexed and sequenced in one lane (HiSeq 2500, Rapid Run Mode, 100 bp paired-end reads) at the University of Minnesota Genomics Center (UMGC) (MN, USA) using Illumina Real Time Analysis software version 1.18.64 for quality-scored base calling.

SMRT reads were assembled using FALCON version 0.7.3 (<u>https://github.com/PacificBiosciences/FALCON-integrate/tree/funzip_052016</u>). After several trial assemblies, a set of parameters was selected with a relatively stringent overlap length to reduce mis-assembly of repetitive regions while maintaining a high contiguity (**Text S1**). The read length cutoff was auto-computed as 9,691 bp for 12NC29 and 8,765 bp for 12SD80. After assembly, FALCON-Unzip (31) was used to phase haplotypes and generate consensus sequences for primary contigs and haplotigs using default parameters. Primary contigs and haplotigs were polished using the Quiver algorithm and corrected for SNPs and indels using Illumina data via Pilon with parameters --diploid and --fix all (58).

Low-quality contigs (over 20% of their size masked by Quiver and smaller than 100 kbp) were removed using custom python scripts. Eleven contigs from 12NC29 and 2 contigs from 12SD80 with significant hits to non-fungal organisms (BLAST search against the NCBI nr/nt database) were excluded as contaminants. Final assembly metrics were derived using QUAST version 4.3 (59) and the Integrative Genomics Viewer (IGV) (60) was used to visualize haplotig regions in primary contigs. To evaluate assembly completeness, the fungal lineage set of orthologs in the software BUSCO (v2.0) (34)was used for comparison, with *Ustilago maydis* as the species selected for AUGUSTUS gene prediction.

### RNA isolation

Seven day-old oat seedlings were inoculated with 10 mg spores/ml or mock-inoculated with oil. Three leaves were pooled per biological replicate at 2 and 5 days post inoculation (dpi), frozen in liquid nitrogen and kept at −80°C. Haustoria were isolated from infected leaves at 5 dpi (inoculated with 20 mg spores/ml) as previously described (18) and stored at −80°C. Prior to RNA extraction, haustorial cells were resuspended in 500 μl of RLT lysis buffer (Qiagen), transferred to FastPrep Lysing beads (MP Biomedicals) and homogenized at 6,000 rpm for 40 seconds using a bead-beating homogenizer. Germinated urediniospores (16 hours) were frozen in liquid nitrogen and kept at −80°C. Three biological replicates were performed for each condition. Samples were ground in liquid nitrogen and RNA was extracted using the RNeasy Plant Mini Kit (Qiagen) according to the manufacturer’s protocols. RNA quality was assessed using an Agilent 2100 Bioanalyzer.

### RNA sequencing and transcriptome assembly

Strand-specific RNA library construction and sequencing (Illumina HiSeq 2500 125 bp PE reads) was carried out at the UMGC. Libraries from germinated spores, *in planta* infections, and mock conditions were multiplexed across three lanes, while libraries from haustoria samples were multiplexed across two lanes. Short-reads and low quality bases were trimmed using Trimmomatic (61) with parameters: ILLUMINACLIP 2:30:10 LEADING 3 TRAILING 3 SLIDINGWINDOW 4:10 and MINLEN 100. *De novo* transcriptome assembly was performed separately for each isolate using combined reads from germinated spores, infected plants and haustoria using Trinity v2.4.0 with parameters: --SS_lib_type RF --normalize_reads (37). The combined reads were also mapped to the assembled genomes of each isolate using HISAT2 v2.0.5 (62) with parameters: --rna-strandness RF --no-mixed. Genome-guided assemblies were generated using Trinity with parameters: --SS_lib_type RF --genome_guided_max_intron 3000 --normalize_reads.

### Genome annotation

Each *Pca* assembly (primary contigs and haplotigs combined) was annotated with Funannotate (version 0.6.0, <u>https://github.com/nextgenusfs/funannotate</u>) in diploid mode using transcript evidence from HISAT2 RNAseq alignments, *de novo* Trinity assemblies, genome-guided Trinity assemblies, and EST clusters from the Department of Energy-Joint Genome Institute (DOE-JGI) for the Pucciniomycotina group (downloaded Feb 20, 2017, <u>http://genome.jgi.doe.gov/pucciniomycotina/pucciniomycotina.info.html</u>). The Funannotate pipeline ran the following: i) repeats were identified using RepeatModeler (63) and soft-masked using RepeatMasker (64), ii) protein evidence from UniProtKB/SwissProt curated database (downloaded on April 26, 2017) was aligned to the genomes using TBLASTN and exonerate (65), iii) transcript evidence was aligned using GMAP (66), iv) *ab initio* gene predictors AUGUSTUS v3.2.3 (67) and GeneMark-ET v4.32 (68) were trained using BRAKER1 (69), v) tRNAs were predicted with tRNAscan-SE (70), vi) consensus protein coding gene models were predicted using EvidenceModeler (71), vii) and finally gene models were discarded if they were more than 90% contained within a repeat masked region and/or identified from a BLASTp search of known transposons against TransposonPSI (72) and Repbase repeat databases (73). Any fatal errors detected by tbl2asn (<u>https://www.ncbi.nlm.nih.gov/genbank/asndisc/</u>) were fixed. Functional annotation used available databases and tools including PFAM (74), InterPro (75), UniProtKB (76), MEROPS(77), CAZymes (78), and a set of transcription factors based on InterProScan domains (79) to assign functional annotations (full list at <u>https://github.com/nextgenusfs/funannotate</u>). Functional annotations for each isolate were compared (compare function) and summary heatmaps prepared from the parsed results using ComplexHeatmap (1.12.0) in R. Gene ontology (GO) terms were compared between isolates using goatools with Fisher’s exact test with false discovery rate and multiple test correction (<u>https://github.com/tanghaibao/goatools</u>).

### Identification of collapsed and haplotig-associated regions, telomeres and GC content analysis

Primary contigs and haplotigs were aligned pair-wise using NUCmer (80) with default parameters. A customized script was used to determine coordinates for matches between primary contigs and haplotigs by scanning aligned blocks along the primary contigs and chaining the aligned haplotig blocks located within 15 kbp. Illumina DNA-sequencing reads were mapped to primary contigs and haplotigs with BWA-MEM version 0.7.12 with default parameters. SAM alignment files were sorted and converted to BAM files with SAMtools (v1.3) (81) and to BED format with BEDtools (v2.25) (82). Coverage was estimated using BEDtools complement and coverage and assigned to genomic regions using the haplotig-region coordinate files. Coverage distributions were plotted as density histograms with the ggjoy package in R. The GC content of all contigs was calculated and the distribution plotted with the hist function in R. Telomeres were identified by the presence of at least 10 repeats of CCCTAA or TTAGGG within 200 bp of the end of a contig using a custom script.

### Genome-wide heterozygosity and variant analysis

Small variants (SNPs and indels) were identified by mapping Illumina DNA-sequencing reads to only the primary contigs of each assembly using BWA-MEM version 0.7.12 with default parameters. PCR duplicates were removed using SAMtools (v1.3) (81) and SNPs were called using FreeBayes (v1.1.0) (83). SNPs were filtered using vcflib (v1.0.0-rc1, <u>https://github.com/vcflib/vcflib</u>) with parameters (QUAL > 20 & QUAL / AO > 10 & SAF > 0 & SAR > 0 & RPR > 1 & RPL > 1 & AB > 0.2 & AB < 0.8) within isolates or (QUAL > 20 & QUAL / AO > 10 & SAF > 0 & SAR > 0 & RPR > 1 & RPL > 1) between isolates. Variants were annotated for genomic location and functional impact using ANNOVAR (2017 Jul 16 version) (84).

*K-mer* counts (21 bp) were generated with Jellyfish (v2.1.3) from raw Illumina DNA sequencing data of *Pca* isolates as well as Illumina sequencing data downloaded from the NCBI SRA for the rust species: *Melampsora larici-populina* (SRR4063847) (17), *Puccinia striiformis* f. sp. *tritici* (SRR058505 and SRR058506) (19), *Puccinia triticina* (SRR027504 and SRR027505), and *Melampsora lini (22)*. The resulting histograms were used as input for GenomeScope (41).

To identify structural variations (SV), haplotigs were aligned to primary contigs with MUMmer (v3.23) with parameters: nucmer -maxmatch −1 100 -c 500 (80). SVs were detected with Assemblytics (42) using default parameters with a minimum variant size of 50 bp, a maximum variant size of 10 kbp, and a unique sequence length for anchor filtering of 10 kbp.

### Identification of alleles and orthologs between isolates

Proteinortho (38) with parameters: -e 1e-05 -synteny -singles was used to identify orthologous groups based on all-against-all blastp search of all annotated genes in 12SD80 and12NC29, followed by construction of an edge-weighted directed graph (edge weight = blast bit score), and heuristic identification of maximal complete multipartite subgraphs. Protein nodes included in subgraphs were defined as orthologous groups. Orthologous genes located in homologous haplotig and primary contig regions based on a gene annotation (gff3) file were assigned as allele pairs.

### Secretome and effector prediction and expression analysis

Secreted proteins were predicted using a method sensitive to fungal effector discovery (85) based on: (i) the presence of a predicted signal peptide using SignalP-NN 3.0 (86), (ii) a TargetP localization prediction of “secreted” or “unknown” (with no restriction on the RC score) (87), and (iii) no transmembrane domain outside the signal peptide region (with TMHMM 2.0) (88). Secreted effectors were predicted using EffectorP 1.0 (44). FeatureCounts (89) was used to generate read counts for each gene from RNAseq data and genes differentially expressed in either haustoria or infected leaves relative to germinated spores (|log fold change| > 1.5 and an adjusted *p*-value < 0.1) were identified using the DESeq2 R package (90). k-means clustering was performed on average rlog transformed values for each gene and condition. The optimal number of clusters was defined using the elbow plot method and circular heatmaps drawn using Circos (91). Gene ontology (GO) enrichment analysis was carried out with the enrichGO function in the R package clusterProfiler version 3.4.4 (92) using the “Molecular function” ontology method and the Holm method to correct *p*-values for multiple comparisons. Local gene density was assessed using the method of Saunders et al. (54), with updates from density-Mapr (<u>https://github.com/Adamtaranto/density-Mapr</u>) to plot the 5’ and 3’ intergenic distance for each gene. The R package GenometriCorr (55) was used to test for associations between effectors and various categories of repeats within 10 kbp regions using default parameters.

### Data and script availability

All raw sequence reads generated and used in this study are available in the NCBI BioProject (PRJNA398546). Genome assemblies and annotations are available for download at the DOE-JGI Mycocosm Portal (http://genome.jgi.doe.gov/PuccoNC29_1 and http://genome.jgi.doe.gov/PuccoSD80_1). Unless specified otherwise all scripts and files are available at <u>https://github.com/figueroalab/Pca-genome</u>.

## Acknowledgments

We thank Dr. Kevin Silverstein at the Minnesota Supercomputing Institute for discussions during genome assembly and analysis. At the USDA-ARS Cereal Disease Laboratory, we thank Roger Caspers for his assistance in maintaining the *Pca* isolates and assigning virulence phenotypes, and Drs. Les Szabo and Jerry Johnson for their assistance during DNA isolation. We also acknowledge Prof. Mark Farman at the University of Kentucky for his input while identifying telomere sequences, and Dr. Matthew Seetin at Pacific Biosciences for assistance with FALCON and FALCON-Unzip.

## Supplemental Material Legends

**Dataset S1-12**. Effector genes on primary contigs and haplotigs with allele pairs for 12SD80 and 12NC29 (Datasets **S1**-**S4**), singleton-effector genes on primary contigs and haplotigs (Datasets **S5**-**S8**), and orthologous and isolate-singleton effectors (Datasets **S9**-**12**). Asterisks in the datasets indicate no ortholog in that genome, and commas between gene and contig names within a genome indicate putative paralogs.

**Text S1**. FALCON config file parameters

**Table S1.** Summary statistics for SMRT sequencing reads

**Table S2.** Alignment statistics of RNAseq reads mapping to *Pca* assemblies (primary contigs)

GS, 2, 5, and H indicate germinated spores, 2 dpi, 5 dpi, and haustoria samples, respectively. R1, R2, and R3 designate the different biological replicates.

**Table S3**. Non-redundant GO terms present in predicted effectors on primary contigs.

**Figure S1**. SMRT sequencing output for two *Pca* isolates

Length distributions of filtered polymerase reads (**A**) and subreads (**B**) for 12SD80 (top) and 12NC29 (bottom).

**Figure S2**. Coverage of 12NC29 and GC content of genome assemblies for each *Pca* isolate

(**A**) Density histograms of mean coverage depth of collapsed and haplotig regions of primary contigs, haplotigs, and primary contigs without haplotigs in 12NC29. (**B**) GC content distribution of contigs from 12NC29 and 12SD80 assemblies.

**Figure S3**. Inter-isolate read mapping coverage of isolate-singleton and orthologous genes

Reads from one isolate were mapped to the other isolate to assess coverage of isolate-singleton and orthologous genes on primary contigs. Density histograms of average coverage depth per gene for 12SD80 (left) and 12NC29 (right).

**Figure S4**. Small sequence variants and structural variation between haplotypes of 12SD80 and 12NC29

(**A**) Genome-wide characterization of SNPs and small indels classified by genomic location as intergenic (dark green), 1 kbp downstream (orange) or upstream of a gene (purple), exonic (red) and intronic (light green) in 12SD80 and 12NC29. (**B**) Structural variation between haplotigs and primary contigs that overlap with annotated genes. Colors indicate different classes of SV (shown in the key). (**C**) Distribution of small variants in and around predicted effectors on primary contigs of 12SD80 and 12NC29. Same key as is shown in (**A**). (**D**) SV types in predicted effector genes as in (**B**).

**Figure S5**. GenomeScope analysis of rust species

Comparison of 21 *k-mer* profiles of 12SD80, 12NC29, *Melampsora larici-populina*, *Puccinia striiformis*, *Puccinia triticina*, *Melampsora lini*. Overall heterozygosity rate estimates are shown in each graph.

**Figure S6**. Intra-isolate structural variants

Graph shows size distribution of structural variants from 50-10,000 bp in haplotigs relative to primary contigs of (**A**) 12SD80 and (**B**) 12NC29 identified using Assemblytics.

**Figure S7**. GO enrichment analysis of secreted proteins

Number of genes in enriched GO term classes in the secreted protein sets of 12SD80 and 12NC29. Dot sizes represent the ratio of a given term out of all enriched GO terms, and colors indicate the adjusted *p*-value according to the scale insets.

**Figure S8**. Secretome clustering and orthology between individual 12SD80 clusters and all 12NC29 clusters

The heatmaps show rlog transformed expression values for germinated spores (GS), isolated haustoria (H), and infected tissues at 2 (2d) and 5 dpi (5d) with dark blue indicating high expression according to the scale inset. Links depict orthology relationships between secretome genes (grey lines) and effectors (red lines) in all 12NC29 clusters, and 12SD80 clusters (**A**) 1, (**B**) 2, (**C**) 3, (**D**) 6 and (**E**) 7.

**Figure S9**. Secretome clustering and orthology between individual 12NC29 clusters and all 12SD80 clusters

The heatmaps show rlog transformed expression values for germinated spores (GS), isolated haustoria (H), and infected tissues at 2 (2d) and 5 dpi (5d) with dark blue indicating high expression according to the scale inset. Links depict orthologous relationships between secretome genes (black lines) and effectors (red lines) in all 12SD80 clusters, and 12NC29 clusters 1 (**A**) 1, (**B**) 2, (**C**) 3, (**D**) 6 and (**E**) 7.

